# An iron acquisition system promotes *Bifidobacterium longum* fitness in competitive early life gut communities

**DOI:** 10.64898/2026.11.15.766743

**Authors:** Luke Acton, Cho Zin Soe, Magdalena Kujawska, Jerry Augustine, Dodzi Kwaku Jnr. Senoo, Hannah Trivett, Shabhonam Caim, Ayuki Shimpo, Callum Clark, Lisa Friess, Douwe van Sinderen, Peter T Chivers, Lindsay J Hall

## Abstract

Iron availability influences the composition of the gut microbiota, yet the mechanisms that enable beneficial bacteria to acquire iron and compete within iron-variable communities remain poorly understood. Here, we identify Fei (<u>Fe</u> import), an iron-responsive acquisition system that promotes the fitness of *Bifidobacterium longum* subsp. *longum*, a prevalent member of the early-life gut microbiota. Comparative profiling of 20 bifidobacterial strains revealed substantial inter- and intraspecies variation in growth under iron limitation, with multiple *B. longum* subsp. *longum* strains maintaining comparatively robust growth. Transcriptomic analysis identified a four-gene locus, *feiMQOX*, that was strongly induced during iron depletion and returned towards control expression following iron supplementation. The complete locus was highly conserved within *B. longum* subsp. *longum* but unevenly distributed across the wider genus. Disruption of individual Fei components impaired growth and reduced cell-associated iron under iron-restricted conditions, while metal-binding studies of purified FeiX demonstrated a physiological preference for Fe(II). FeiX also provided a pronounced fitness advantage during competition with an infant-derived *Enterococcus faecalis* isolate, particularly under iron limitation. Finally, analysis of longitudinal metagenomic data from Bangladeshi infants demonstrated that Fei is present within infant gut communities and that its abundance changes following iron supplementation. Together, these findings identify a lineage-associated iron-acquisition system that connects micronutrient availability with bifidobacterial physiology and competitive fitness. More broadly, our findings establish strain-level micronutrient acquisition as a potentially important determinant of beneficial bacterial fitness within the developing gut microbiota.

## Introduction

Iron is an essential micronutrient that supports fundamental cellular processes across all domains of life. In bacteria, iron acts as a cofactor in enzymes involved in central metabolism, DNA synthesis, and protection from oxidative stress (1–4) . However, the sensitivity of ferrous ion to oxidation and the limited solubility of Fe(III) severely restrict iron bioavailability within host-associated environments. In the gastrointestinal tract, iron accessibility is further shaped by local pH, dietary and microbial ligands, host sequestration mechanisms and competition between microorganisms (5). Gut microbes must therefore acquire iron from a limited and dynamically changing pool, making iron acquisition a potentially important determinant of microbial fitness and community assembly.

These ecological pressures are particularly important during early life, when the gut microbiota undergoes rapid development alongside major changes in diet, host physiology and micronutrient availability (6). Iron requirements are high during infancy, and iron deficiency anaemia (IDA) remains common among infants and young children worldwide (7). Oral iron supplements and iron-containing micronutrient powders are consequently used widely to prevent or treat iron deficiency. However, only a proportion of orally administered iron is absorbed in the upper gastrointestinal tract, with unabsorbed iron reaching the colon and becoming accessible to the resident microbiota (8). Iron interventions have been associated with altered microbiota compositions in some settings, including reductions in beneficial commensals and expansion of potentially pathogenic bacteria, although the magnitude and direction of these effects vary between cohorts, host iron states and treatment contexts. For example, a recent placebo-controlled trial involving ∼900 Bangladeshi infants identified treatment-associated trends towards reductions in commensal taxa and increases in potential pathogens, particularly among iron-replete infants (9). Together, these findings indicate that microbial responses to iron interventions are heterogeneous and may depend on the iron-acquisition capabilities of individual community members.

Most mechanistic understanding of bacterial iron acquisition derives from pathogens, which use siderophores, ferrous and ferric iron transporters, and receptors for host-associated iron complexes to support growth and colonisation (10). Members of the Enterobacteriaceae, for example, possess highly efficient iron-scavenging systems that can provide a competitive advantage within the inflamed or iron-supplemented gut (11). Far less is known about iron acquisition by beneficial anaerobic members of the gut microbiota acquire iron or compete with other microorganisms when iron availability changes. This represents an important knowledge gap because micronutrient acquisition may influence not only their own persistence, but also their ability to compete with opportunistic competitors.

Bifidobacteria are dominant members of the early life gut microbiota and are particularly abundant in healthy breast-fed infants (12). Through their metabolism of human milk oligosaccharides and other host- and diet-derived carbohydrates, bifidobacteria contribute to microbial community structure, intestinal barrier function and colonisation resistance (13–15). However, the abundance and species composition vary considerably among infants and can change during nutritional or ecological perturbation (16). Although carbohydrate utilisation by *Bifidobacterium* spp. has been studied extensively, considerably less is known about the micronutrient-acquisition systems that allow particular species and strains to establish and compete within the developing gut microbiota.

Available evidence indicates that iron-acquisition capacity is not uniform across the genus. The ferrous iron transporter *bfeUOB*, has been characterised in *Bifidobacterium* breve UCC2003, although disruption of the system did not significantly impair gastrointestinal colonisation in the models tested (17,18). Proteomic and genomic analyses have identified additional putative iron-acquisition proteins in other bifidobacterial species, but the substrates, regulation and ecological functions remain unresolved (19). It is therefore unclear whether bifidobacterial lineages possess specialised iron-acquisition mechanisms that allow them to withstand iron limitation and maintain fitness during competition with other members of the gut microbiota.

Here, we identify Fei, a previously uncharacterised, iron-responsive acquisition system in *B. longum* subsp. longum. The *feiMQOX* locus that was induced by iron depletion, conserved across *B. longum* subsp. *longum* strains, supports iron accumulation and growth under iron restriction. Fei also promoted competitive fitness against *Enterococcus faecalis*, while analysis of a Bangladeshi infant cohort showed *Fei* abundance changed following iron supplementation. Together, these findings identify a lineage-associated mechanism linking iron acquisition to bifidobacterial fitness in the developing gut microbiota.

## Materials and Methods

### Bacterial culture conditions

Bacterial strains and primers used in this study are listed in Supplementary Table 1 and 2. *Bifidobacterium longum* subsp. *longum* LH_277 was isolated from a stool sample collected from a healthy, human-milk fed, 11-week-old infant in accordance with protocols approved by the National Research Ethics Service (NRES)-approved UEA/QIB Biorepository (Licence no. 11208) and the Quadram Institute Bioscience Ethics Committee.

Unless otherwise stated, *Bifidobacterium* strains were cultured anaerobically at 37°C in Reinforced Clostridial Medium or on Reinforced Clostridial Agar (Oxoid) supplemented with 0.06% (w/v) L-cysteine. Anaerobic conditions were maintained under an atmosphere containing 80% N, 10% CO, 10% H . For solid media agar was added to a final concentration of 1.5% (w/v).

Iron-restrictive medium (IRM) was prepared as described previously by Petry *et al.*, with modifications (20). The basal medium contained 5.64 g/L 5× M9 salts, 3 g/L peptone, 1 g/L yeast extract, and 11.1 mM D-glucose. Following autoclaving, sterile-filtered L-cysteine 0.02% (w/v), MgSO (final concentration 0.40 mM), and CaCl (final concentration 20 μM) were added aseptically to the cooled medium. Where indicated, 2,2′-dipyridyl (DP) was added from a sterile-filtered 50 mM stock solution to further restrict bioavailable iron.

### Growth of *Bifidobacterium* strains under iron restrictive conditions

To assess bifidobacterial growth under iron-depleted conditions, strains were cultured anaerobically in RCM for 48 h. Cultures were centrifuged at 3,000 × *g* for 5 mins, and cell pellets were washed three times with sterile phosphate-buffered saline (PBS) before resuspension in 10 mL PBS.

Washed cells were used to inoculate 200 µL cultures of RCM, IRM) or, IRM supplemented with 250 µM DP. Cultures were inoculated to an initial optical density (OD) of 0.01 and incubated under the conditions described above for 24 h. Growth was monitored by measuring OD every 15 min using a Cerillo Stratus microplate reader 600nm (Cerillo, USA). Experiments were performed using 2 independent biological replicates.

### RNA extraction and identification of LH_277 iron-responsive genes

To identify iron-responsive genes, overnight LH_277 cultures grown in de Man, Rogosa and Sharpe broth (MRS;Difco) were used to inoculate three experimental conditions in biological triplicate: control (IRM), iron-depleted (IRM + 100 µM DP), and iron-supplemented (IRM + 100 µM DP supplemented with 50 µM FeCl). Cultures were harvested during lag phase (∼6 h), early exponential phase (∼14 h), and stationary phase (∼24 h). Cells were washed with sterile PBS and RNA extraction was performed immediately. Cell pellets were resuspended in 600 µL RLT lysis buffer containing 8 µL β-mercaptoethanol and transferred to Lysing Matrix E tubes (MP Biomedicals). Cells were disrupted using a FastPrep-24 instrument (MP Biomedicals) at 6.0 m/s for three 1-min cycles, with samples incubated on ice for 5 min between cycles. Lysates were centrifuged at 14,000 × *g* for 10 min, and total RNA was extracted from the supernatant using the RNeasy Mini Plus kit (Qiagen) according to the manufacturer’s instructions.

RNA concentration and integrity were assessed using an Agilent 2100 Bioanalyzer (Agilent Technologies). Samples with RNA integrity number (RIN) values ≥8 were subjected to ribosomal RNA depletion using Illumina Ribo-Zero Plus rRNA Depletion kit. RNA samples were submitted to the Wellcome Sanger Institute (UK) where libraries were prepared using Illumina TruSeq RNA Library prep kit and sequenced on an Illumina HiSeq V4 platform using a 75 bp paired-end, non-stranded sequencing.

## Differential expression analysis

Raw sequencing reads were quality assessed and trimmed using Trim Galore! (version 0.4.2) and default parameters. Following quality filtering, residual ribosomal RNA reads were removed using SortMeRNA (version 2.1b). Filtered, reads were pseudoaligned to the LH_277 genome assembly 19109_5#24 using Kallisto (version 0.43.1).

Differential expression analysis was performed in R using Hall laboratory scripts (https://github.com/Shabhonam/RNAseq) and the DESeq2 package. Each growth phase was analysed independently, and comparisions were made between control and Iron deplete or Iron replete conditions. Genes with a Benjamini-Hochberg-adjusted *p*-value (padj) < 0.05 and an absolute log_2_ fold change of at least 0.5 were considered significantly differentially expressed.

### Insertional mutagenesis in *Bifidobacterium longum* subsp. *longum*

Insertional mutants were generated in *Bifidobacterium longum* NCIMB 8809, a strain for which genetic manipulation has previously been established (21). Internal fragments of *feiM, feiQ, feiO* and *feiX* were amplified from NCIMB8809 genomic DNA using primers listed in Supplementary Table 2. PCR products were ligated into pFREM2 using *Xba*I and *Nco*I restriction sites for *feiMQX*, and *Xba*I and *Hind*III restriction sites for *feiO*.

Recombinant plasmids were introduced into *Escherichia coli* EC101 by electroporation (1.8 kV, 20 µF, 200 Ω). Following a 90-minute recovery period, cells were plated onto Luria– Bertani (LB) agar supplemented with 250 µg/mL erythromycin. Successfully ligated plasmids were confirmed by PCR, using primers flanking the insertion region before transformation into NCIMB 8809.

Electrocompetent NCIMB 8809 cells were prepared as previously described, with minor modifications (22). A single colony was cultured in MRS broth at 37°C for 16 h and subsequently subcultured twice in MRS broth supplemented with 7% (w/v) sucrose and 122.3 µM iron(II) sulfate heptahydrate. Cultures were grown to an OD_600_ of 0.9–1.1 and used to inoculate modified MRS broth (10% inoculum in a 50 mL final volume). Modified MRS was prepared from individual components and supplemented with 1% lactose, 7% sucrose, and 20 mM NaCl. Cells were harvested before the OD600 exceeded 0.5.

Cells were centrifuged (3000 × g for 10 minutes) and washed three times with 1 mL sucrose-citrate buffer containing 0.5 M sucrose, 1 mM citrate. Washed cells were resuspended in 200 µL sucrose-citrate buffer. All culture steps were performed under anaerobic conditions.

Electrocompetent cells were combined with between 0.5-1 µg plasmid DNA and incubated on ice for 30 min in a 2 mm electroporation cuvette. Cells were electroporated at 2.5 kV, 20 µF, 300 Ω. Following electroporation, cells were recovered for 3 h at 39°C in modified MRS containing 0.90 µM CaCl and 0.245 µM 1,4-dihydroxy-2-naphthoic acid (DHNA). Transformants were selected on RCA containing 0.90 µM CaCl, 0.245 µM DHNA, and 100 µg/mL erythromycin and incubated anaerobically for 2–4 days. Insertional disruption was confirmed by long read whole-genome sequenced performed by MicrobesNG, UK. Genomes were assembled using Flye v2.9.2 (23) and annotated using Bakta v1.9.3 (24). Gene disruption was confirmed via manual inspection of the FeiMQOX locus using Artemis (25)

## Growth characterisation of Fei mutants

Parental NCIMB 8809 and Fei insertional mutants were cultured anaerobically in RCM for 48 h. Cultures were washed three times with sterile PBS and used to inoculate RCM, IRM, IRM containing 200 µM 2,2′-dipyridyl, or IRM containing 200 µM 2,2′-dipyridyl and 75 µM ferric nitrate at a starting OD of 0.05.

For liquid growth assays, 200 µL cultures were incubated anaerobically at 37°C, and OD was measured every 15 min for 24 h. For spot-dilution assays, cultures were adjusted to an OD of 1, serially diluted ten-fold and spotted onto the indicated agar media. Plates were incubated anaerobically at 37°C for 48 h before imaging. Experiments were performed using three independent biological replicates.

## Phylogenetic analysis and identification of Fei components

Genome assemblies and predicted amino acid sequences from publicly available and in-house *Bifidobacterium* strains were used to examine the distribution of Fei components. Publicly available assemblies were downloaded from NCBI, while genomes from in-house strains (LH_X) were generated and processed as described previously (26).

Taxonomic classification and marker-gene alignment were performed using GTDB-Tk v2.3.2 with the classify-wf workflow. The resulting alignment was used to infer a maximum-likelihood phylogeny in IQ-TREE v2.3.2 (27) using best-fit model (LG+F+I+R7). The phylogeny was visualised iTOL v7 (28). Gain and loss events were inferred using Dollo parsimony in Count v10.04 (29).

Fei components were identified by comparing amino acid sequences encoded by the LH_277 *feiMQOX* locus against the predicted bifidobacterial protein sequences using local blastp (e-value < 1e−5, identity and coverage 30%).

## Metal content determination of whole cells and FeiX

Parental NCIMB 8809 and Fei mutant strains were grown anaerobically for 18 h at 37°C in 12 mL MRS supplemented with 0.07% (w/v) L-cysteine. Cultures were diluted into 12 mL IRM supplemented with 0.07% (w/v) L-cysteine to a starting OD_600_ of 0.010 and incubated for a further 24 h at 37°C.

Following incubation, OD_600_ values were measured. Aliquots were serially diluted, and 100 µL were plated on MRS agar supplemented with 0.07% L-cysteine to determine CFU/mL. The remaining 11 mL was centrifuged at 4 °C, 15 min, 3900 x g.

Cell pellets were washed four times in 10 mL ice-cold buffer containing 10 mM HEPES, pH 7.5, 0.5 M sorbitol, and 0.1 mM EDTA. Washed pellets were digested in 500 µL of Ultrapure HNO_3_ 65% (v/v) for 48 h.

Elemental analysis was performed at the Durham University Bio-ICP-MS Facility using Thermofisher iCAP RQ. Calibration standards were prepared in HNO_3_ 65% (v/v) to match the sample matrix, with silver, beryllium, and indium used as internal standards. Calibration standards and digested samples were diluted 10-fold into 2.5% nitric acid. Iron content was normalised to CFU count and expressed as iron atoms per CFU. Metal content of FeiX samples was determined in a similar manner to whole cells. However, calibration standards contained 20 mM Hepes, pH 7.5 and 200 mM NaCl instead of HNO_3_ 65% (v/v) to match the protein sample matrix.

## FeiX expression vector construction

A codon-optimised *B. longum feiX* gene was synthesised by Eurofins. The gene was excised from the manufacturer provided plasmid using *Nde*I and *Xho*I and ligated into pET21b digested with the same restriction enzymes to generate pET-*feiX*. The construct was introduced into *E. coli* JM109 for plasmid propagation and subsequently transformed into *E. coli* DL41(DE3) for protein expression (30).

Small scale expression tests were performed in 2 mL LB broth containing 100 µg/ml ampicillin. Protein expression was induced with 0.4 mM isopropyl-β-D-thiogalactopyranoside (IPTG) and expression was assessed by SDS-PAGE.

## FeiX expression and purification

*E. coli* DL41(DE3) containing pET*-feiX* was cultured at 37 °C with shaking at 150 rpm in 1 L of LB broth containing 100 µg/mL ampicillin. At OD_600_ ∼0.6, protein expression was induced with 1 mM IPTG and cultures were incubated at 18°C for 18 h.

Cells were harvested by centrifugation at 4°C for 20 min at 3900 x g and resuspended in 30 mL buffer containing 20 mM HEPES, pH 7.5, 100 mM NaCl, 1mM PMSF, and 1 mM EDTA. Cells were disrupted by sonication for 5 min (4 × 10 s on ice, 1 min intervals), then lysates were centrifuged at 4°C for 30 min at 33000 x g. The resulting supernatant was passed through a 0.22 µm filter.

The clarified lysate was loaded at 1 mL/min onto a 5mL HiTrap Q HP column equilibrated with 20 mM HEPES, pH 7.5, 100 mM NaCl, 1mM Tris (2-carboxyethyl)phosphine (TCEP), and 1 mM EDTA. Protein was eluted using a 30 mL gradient to 1 M NaCl. Fractions (1.5 mL) containing FeiX were idenitfed using SDS-PAGE, pooled, and loaded onto a HiLoad 16/600 Superdex 200 pg column equilibrated with 20 mM HEPES, pH 7.5, 200 mM NaCl, 1mM TCEP, and 1 mM EDTA.

Fractions (1.5 mL) containing purified FeiX were pooled and loaded onto a 1 mL HiTrap Q HP column equilibrated with 20 mM HEPES, pH 7.5, and 100 mM NaCl. The column was transferred to an anaerobic globe box (Belle Technology) and protein eluted in buffer containing 20 mM HEPES, pH 7.5, and 1M NaCl. The purified protein was stored at 4°C. Protein concentration was determined from absorbance at 280 nm using a calculated molar extinction coefficient of 52940 M^–1^ cm^–1^ (31), and was in the range of 400 -600 µM.

## Preparation of anaerobic metal stocks

Ammonium iron(II) sulfate hexahydrate was dissolved under anaerobic conditions in N_2_-purged 0.1% (v/v) HCl. The iron oxidation state was confirmed using a 10-fold molar excess of ferrozine and an extinction coefficient of 27,900 M^−1^ cm^−1^ at 562nm. Dilutions were prepared in N_2_-purged ultrapure water. ZnSO_4_ was dissolved in N_2_-purged ultrapure water. Fe(II) and Zn(II) stock concentrations were determined by ICP-MS

## Determination of FeiX metal stoichiometry

Metal-binding stoichiometry was assessed anaerobically in 20 mM Hepes, pH 7.5, and 200 mM NaCl. FeiX at 20 µM was incubated for 1 h with 22 µM Zn (II) or 44 µM Fe(II). Samples were applied to PD-10 desalting columns (Cytiva). Columns were first washed with 1 mL of 5 mM EDTA, to remove any residual metal ions, followed by equilibration with 25 mL assay buffer before sample loading (500 µL), then elution with 8 mL of buffer. Elution fractions (0.5 mL) were collected and metal and protein content determined by ICP-MS and absorbance (A_280_), respectively.

## Determination of FeiX metal affinities

Fe(II)- and Zn(II)-binding affinities were determined by competition with mag fura-2 (MF-2) under anaerobic conditions (ε363 nm=22 000 M^-1^ cm^-1^) (32). Experiments were performed in 20 mM Hepes, pH 7.5 and 200 mM NaCl. Fe(II) or Zn(II) was titrated into a mixture containing FeiX and MF-2, and absorbance was measured at equilibrium at 363 nm.

Data were analysed using DYNAFIT (33). Fe(II) data were fitted to a model describing competition between 1:1 Fe(II):MF-2 with 1:1 Fe(II):FeiX. MF-2 *K*_d Fe(II)_ = 5.3 x 10^-6^ M (32). The FeiX *K*_d Zn(II)_ was determined by fitting the data to a model describing the competition between 1:1 Zn(II):MF-2 with 2:1 Zn(II):FeiX. MF-2 *K*_d Zn(II)_ = 2.0 x 10^-8^ M (32). Confidence intervals were determined using the DYNAFIT Monte-simulation function.

### Bifidobacterium-Enterococcus competition assays

Competition assays were performed using *Enterococcus faecalis* BHI91, isolated from a healthy infant participating in the PEARL cohort (34), and parental NCIMB 8809 or the corresponding *feiX* insertional mutant. To enable selective enumeration, the NCIMB 8809 strains were transformed with pVG-GFP, which encodes spectinomycin-resistance (35).

*E. faecalis* was cultured aerobically overnight at 37°C with shaking at 200 rpm in brain heart infusion (BHI) broth (Sigma-Aldrich). *B. longum* strains were cultured anaerobically overnight as described above. Cultures were adjusted to an OD_600_ of 0.05 and mixed at equal volumes. The resulting suspension were used to inoculate 10 mL RCM, iron-depleted medium or iron-supplemented medium at a combined starting OD_600_ of 0.1. Iron-depleted medium contained basal IRM with 200 µM 2,2′-dipyridyl, while iron-supplemented medium contained 200 µM 2,2′-dipyridyl and 100 µM ferric nitrate.

Co-cultures were incubated anaerobically at 37°C and sampled at 0, 6 and 24 h. Samples were serially diluted ten-fold in PBS. Viable *E. faecalis* was enumerated on RCM incubated aerobically, while *B. longum* was selectively enumerated on RCA supplemented with 200 µg/mL spectinomycin and incubated anaerobically. The proportion of each organism in the recoverable culturable population was calculated from the corresponding CFU/mL values.

## Quantification of Fei in the BRISC cohort

Publicly available metagenomic data from Bangladeshi BRISC cohort was reanalysed to investigate prevalence and abundance of the *feiMQOX* locus following iron supplementation (9). A total of 225 metagenomes were downloaded from NCBI under accession PRJNA1081952. Samples from participants with documented antibiotic exposure were excluded in accordance with the original study metadata.

Reads were trimmed and quality filtered using fastp (v1.3.3) (36). Human-derived reads were removed using Hostile v2.0.0 with default settings (37). Paired baseline and post- intervention samples from 36 infants receiving 12.5mg of elemental iron via ferrous sulphate syrup were selected for locus-level analysis.

Reads were mapped to the *feiMQOX* locus using Bwa-mem2 v2.2.1 (38), mapped reads were counted and processed using SAMtools v1.22.1 (39). Gene abundance was normalised as reads per kilobase per million mapped reads. Microbial composition profiles were generated using Kraken2 v2.1.3 (40) with the standard database downloaded on 31^st^ December 2024, and species-level abundance was estimated using Bracken v2.9 (41).

Statistical analyses were performed in R using tidyverse v1.3.2 (42), vegan v2.7-3 (43), lme4 v2.0-1 (44), lmerTest v3.2-1 (45), emmeans v2.0.2 (46). The final model included time point as a fixed effect and participant identity as a random intercept.

## Metagenome assembly and genome reconstruction

Quality-filtered, host-depleted reads were assembled individually using megahit v1.2.9 (47). Metagenome binning was performed using MetaWRAP v1.3 (47) with CONCOCT v1.1.0 (48), MetaBAT2 v2.12.1 (49), and MaxBin2 v2.2.6 (50)). Bins generated using the three approaches were consolidated using the MetaWRAP bin-refinement module.

Bin quality was assessed using CheckM v1.1.3 (51). Bins with greater than 70% estimated completeness and less than 5% estimated contamination were retained. Taxonomic classification was performed using GTDB-Tk v2.7.2 classify_wf module (52).

Bins classified as *B. longum* were annotated using bakta v1.11.4 (24). Phylogenetic analysis was performed using IQ-TREE v2.4.0 as described above. Predicted amino acid sequences were screened for Fei components using the same blastp approach applied to the isolate genomes.

## Statistical analyses

Statistical analyses were performed using R version 4.6.2. The number of biological replicates is reported in the corresponding figure legends. Data are presented as mean average, with error bars representing standard error.

Growth, ICP-MS and competition data were analysed using a paired t-test. CFU/mL values were log transformed before analysis. For the BRISC analysis, linear mixed-effects models were carried out as described previously. All statistical tests were two-sided, and adjusted *P* values below 0.05 were considered statistically significant.

## Results

### *Bifidobacterium longum* subsp. *longum* exhibits enhanced growth under iron limitation

To determine how iron availability influences bifidobacterial growth, we compared 20 human- and animal-associated *Bifidobacterium* strains in nutrient-rich RCM, iron-restrictive medium (IRM), and IRM supplemented with iron chelator 2,2′-dipyridyl (IRM-DP). Growth responses differed markedly across the strains tested (Figure 1). Several strains maintained robust growth in IRM, whereas others displayed delayed or substantially reduced growth rate relative to RCM. These differences became more pronounced following addition of 2,2′- dipyridyl.

**Figure 1.**
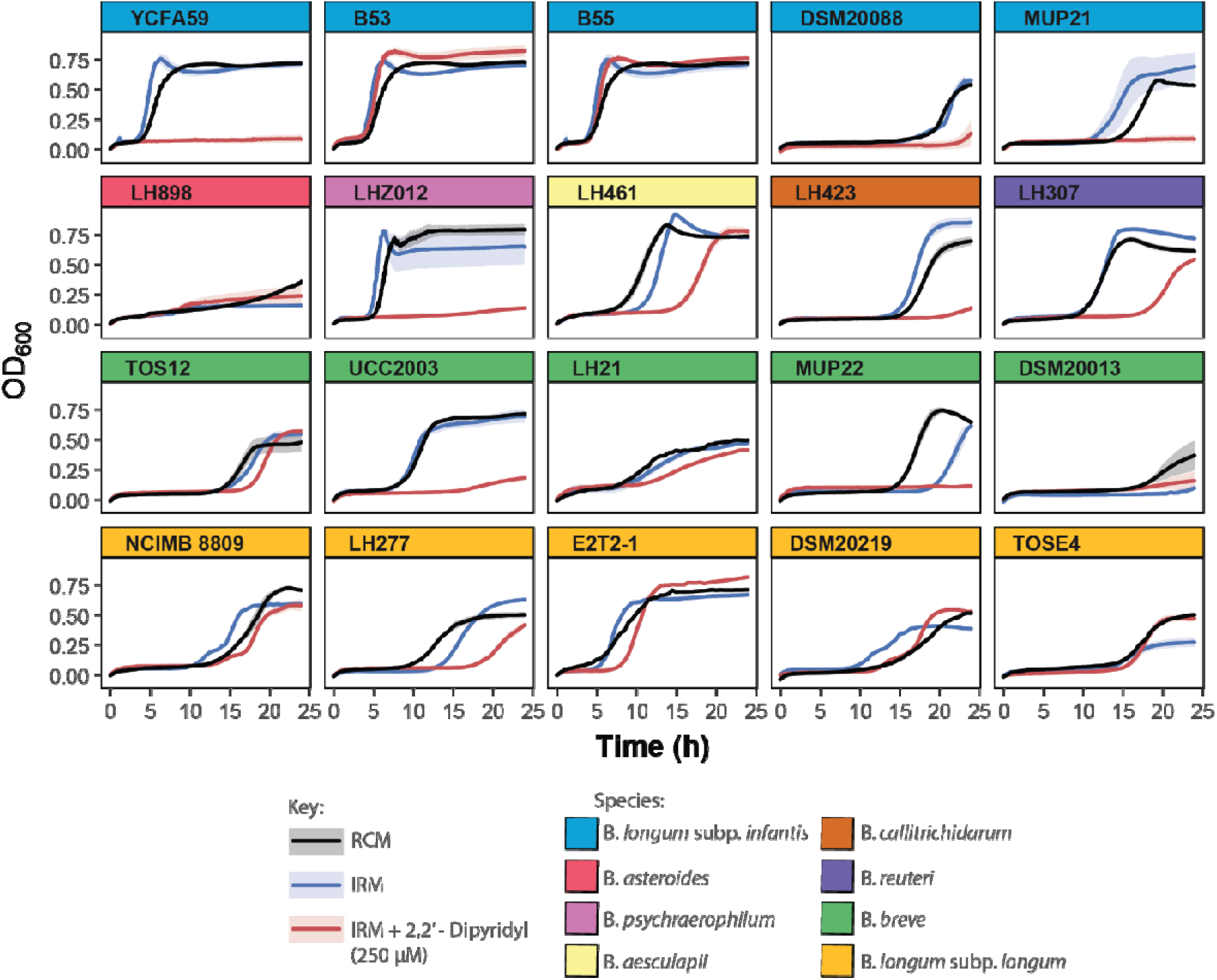
Growth responses of diverse bifidobacterial strains to increasing iron limitation. Twenty *Bifidobacterium* strains representing seven species were cultured in rich culture medium (RCM; black), iron-restrictive medium (IRM; blue), or IRM supplemented with 250 μM 2,2′-dipyridyl (IRM-DP; red), and growth was monitored by OD measurements every 15 min over 24 h. Individual panels show growth curves for each strain, with panel colours denoting species. Shaded areas indicate standard error from two biological replicates. Increasing iron restriction differentially affected growth kinetics, revealing substantial inter- and intraspecies variation in tolerance to iron depletion.

Strains belonging to *Bifidobacterium longum* subsp. *longum*, including NCIMB 8809, E2T2-1, DSM20219, and TOSE4, generally retained growth under iron-restrictive conditions. In contrast, growth of *Bifidobacterium longum* subsp. *infantis* strains YCFA59 and MUP21, and *Bifidobacterium breve* MUP22, were strongly attenuated in IRM-DP. However, responses also varied among strains assigned to the same species or subspecies, indicating that tolerance of iron limitation is not determined solely by taxonomic identity. Collectively, these findings reveal substantial inter-and intraspecies variation in bifidobacterial growth under iron restriction and identify *B. longum* subsp. *longum* as a lineage with comparatively robust growth under the conditions tested.

## Iron limitation induces a candidate Fei acquisition system in *B. longum* subsp

### longum

To identify pathways associated with growth during iron limitation, we profiled the transcriptome of *B. longum* subsp. *longum* LH_277 under control, iron-depleted and iron-supplemented conditions at lag, exponential and stationary phases (Figure 2a). The transcriptional response to iron depletion was strongly growth-phase dependent, with the largest number of differentially expressed genes detected during stationary phase. In total, 380 genes were differentially expressed during stationary phase under iron-depleted conditions relative to control, of which 218 were upregulated and 162 were downregulated.

**Figure 2.**
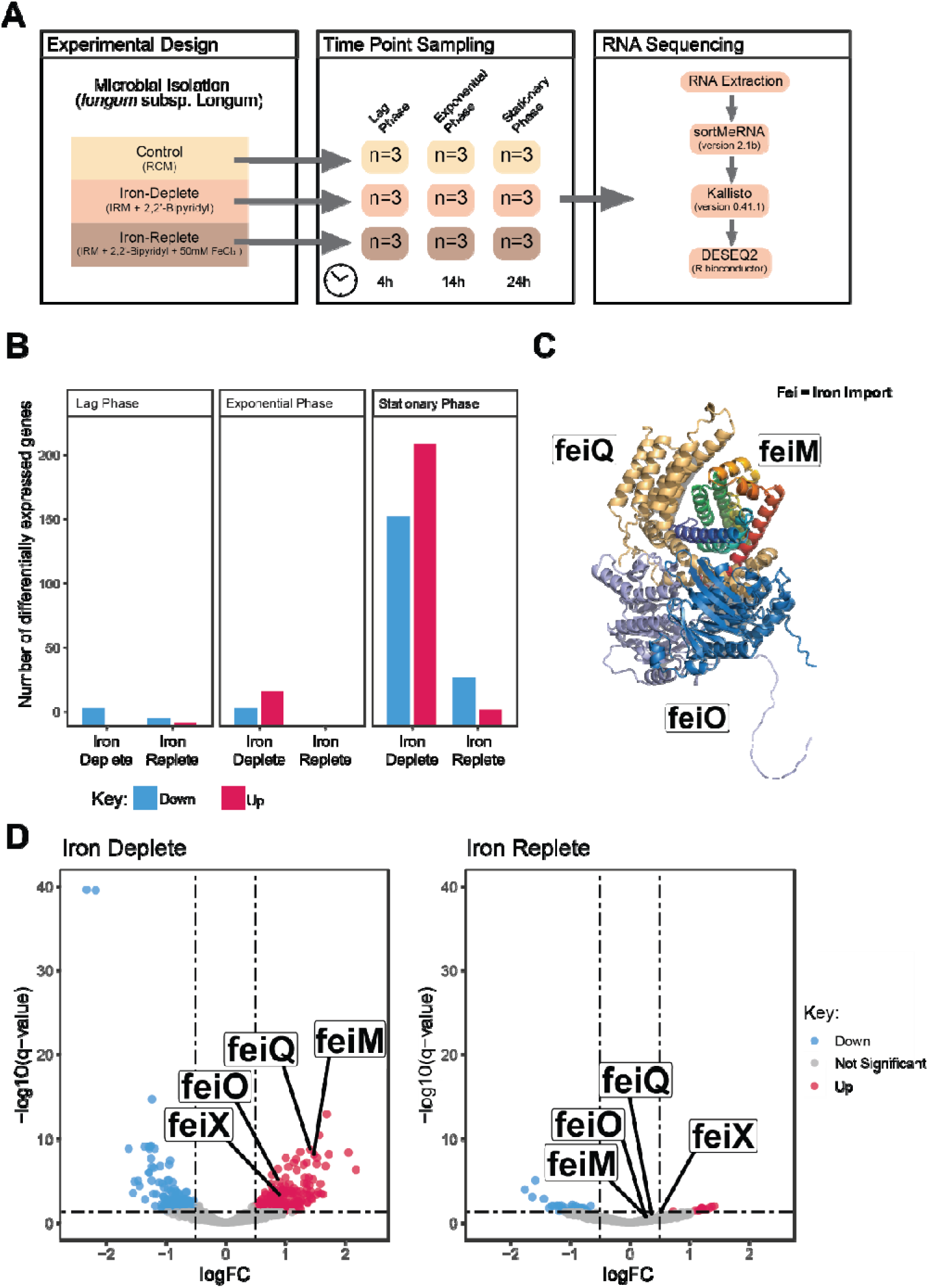
a-d. **Transcriptomic profiling of *Bifidobacterium longum* subsp, *longum* strain LH277. A)** Experimental design and bioinformatic analysis pipeline of RNA sequencing experiment under differential growth phases and iron conditions. B) Number of differentially expressed genes across different growth stages in Iron deplete (IRM + 2’2-Dipyridyl vs control) and Iron replete (IRM + 2’2-Dipyridyl + FeCl_3_ vs control). C) Venn diagrams of overlapping significantly up-or down-regulated genes across growth phases. D) Volcano plots of log fold changes in expression against statistical significance (-log10(q-value)) in stationary phases

Supplementation of iron-depleted cultures with ferric chloride substantially attenuated this transcriptional response, with 96.3**%** of genes induced under iron depletion no longer differentially expressed relative to the control (Figure 2B). This reversal was consistent with the response being driven primarily by iron availability rather than by non-specific effects of 2,2′-dipyridyl.

Among the most strongly induced genes was a four-gene locus encoding proteins with homology to components of an energy-coupling factor transport system (Figure 2D). The genes, subsequently designated *feiM, feiQ, feiO* and *feiX*, were upregulated by 1.52, 1.48, 0.99 and 0.98 log_2_fold changes during stationary phase under iron depletion and returned towards control expression levels following iron supplementation.

Sequence and structural analyses predicted that FeiM, FeiQ and FeiO form membrane-associated components of an ECF-like transporter complex. The locus additionally encoded FeiX, a protein not typically associated with canonical ECF transport systems. AlphaFold modelling revealed structural homology to ZinT, a *E. coli* periplasmic Zn-binding protein. However, FeiX lacks a signal or transmembrane spanning sequence, thus predicting a soluble, cytosolic protein with a function distinct from the additional components of Ni and Co ECF importers (53). Together, the strong induction of the complete locus during iron restriction, reversal following iron supplementation and predicted transporter organisation identified *feiMQOX* as a candidate iron acquisition system for subsequent functional characterisation.

### The complete *feiMQOX* locus is conserved in *B. longum* subsp. *longum* but sporadically distributed across *Bifidobacterium*

To examine the evolutionary distribution of the *Fei* system, we constructed a maximum-likelihood phylogeny incorporating 212 publicly available and 127 in-house *Bifidobacterium* genomes representing 98 species and subspecies. Genome assemblies were classified using GTDB-Tk, and the presence of each Fei component was determined by protein-level sequence comparison with the LH_277 *feiMQOX* locus (Figure 3).

**Figure 3.**
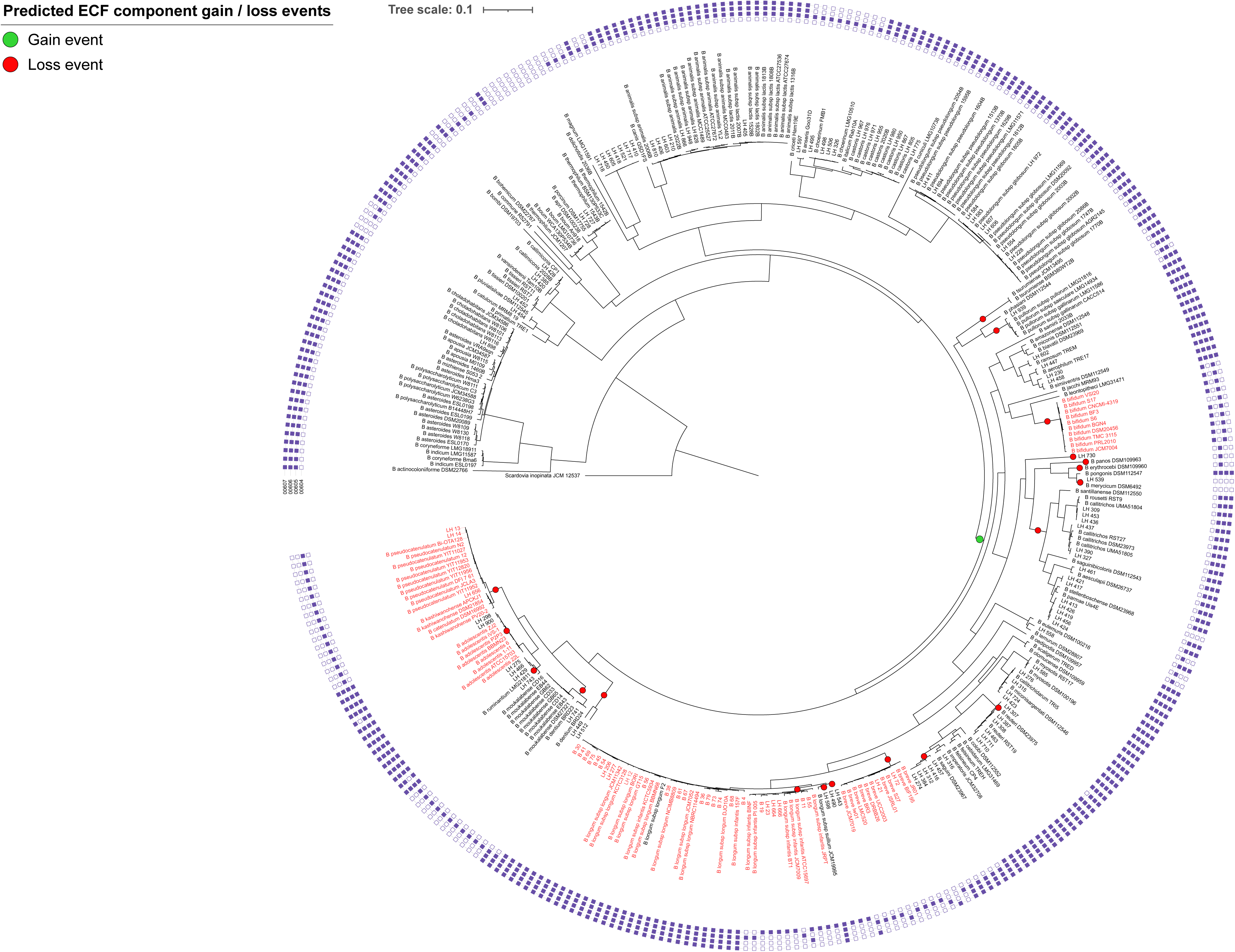
– **Distribution of *FeiMQOX* in *Bifidobacterium* genomes**. Maximum likelihood phylogeny of 339 *Bifidobacterium* strains representing 98 bifidobacterial species and subspecies, constructed using the best-fit model (LG+F+I+R7). Human-associated *Bifidobacterium* strains are marked in red. The genome of *Scardovia inopinata* JCM 12537 was used as an outgroup. The presence of individual Fei iron uptake system components is marked next to the strain identifier (full square = present, empty square = absent). Fei component gain / loss events are mapped onto the tree branches in green and red, respectively. FeiM = 607, FeiQ=606, FeiO=605 and FeiX=604.

The complete four-gene locus was highly conserved within *B. longum* subsp. *longum*. All 28 analysed genomes encoded FeiM, FeiQ, FeiO and FeiX, with high amino acid sequence identity relative to LH_277. Conservation of both gene content and local synteny indicated that the complete system is a stable feature of the *B. longum* subsp. *longum* lineage.

Despite the close evolutionary relationship between the two subspecies, the distribution of Fei components differed markedly in *B. longum* subsp. *infantis*. Only 2 of the 14 *B. longum* subsp. *infantis* genomes analysed contained the complete *feiMQOX* locus. Instead, these genomes typically retained only a few components, although only 3 of the 14 genomes did not encode *FeiX*. This pattern demonstrates that the complete system is not uniformly conserved across the wider *B. longum* lineage.

Outside *B. longum*, the complete locus was identified in 9 out of 10 *B. adolescentis* genomes and in 46 additional genomes assigned to 29 species. Among strains lacking the complete locus, feiX was the gene most frequently absent. Partial loci were more widely distributed across the genus, with individual Fei components detected in all but 12 genomes representing. FeiX was the component most frequently absent from otherwise complete or partial loci, including in strains assigned to *B. animalis* subsp. *animalis*, *B. animalis* subsp. *lactis* and *B. pseudolongum*.

Ancestral-state reconstruction using Dollo parsimony 19 losses and 1 gain of *feiX,* including multiple losses within the *B. longum* subsp. *infantis* lineage (Figure 3). Although these reconstructions do not establish the selective pressures driving locus retention or loss, the strong conservation of the complete system within *B. longum* subsp. *longum* is consistent with a persistent lineage-associated functional advantage. Together, these analyses reveal that Fei is highly conserved within *B. longum* subsp. *longum* but and has uneven evolutionary distribution across the wider genus.

## Fei supports intracellular iron accumulation and growth during iron limitation

To determine whether the Fei system contributes directly to growth under iron limitation, we disrupted each component of the *feiMQOX* locus in *B. longum* subsp. *longum* NCIMB 8809. This strain was selected because reproducible genetic manipulation was previously established in this background (18,22). The *feiMQOX* loci of NCIMB 8809 and LH_277 shared 100% amino acid identity across FeiMQX and 90% between FeiO, with conserved gene order and predicted organisation (Supplementary Figure 1).

Disruption of *feiM, feiQ, feiO* or *feiX* did not produce a substantial growth defect in nutrient-rich RCM or in IRM without additional chelator (Figure 4A and 4B). In contrast, supplementation of IRM with 200 μM 2,2′-dipyridyl revealed a pronounced Fei-dependent phenotype. All four mutants showed significantly reduced growth relative to the parental strain, as demonstrated by spot dilution assays and liquid growth measurements (Figure 4A-C). After 24 h, area under the curve values of the *feiM, feiQ, feiO* and *feiX* mutants were reduced by 40.7%, 41.1%, 38.9% and 19.8%, respectively, relative to the parental strain.

**Figure 4.**
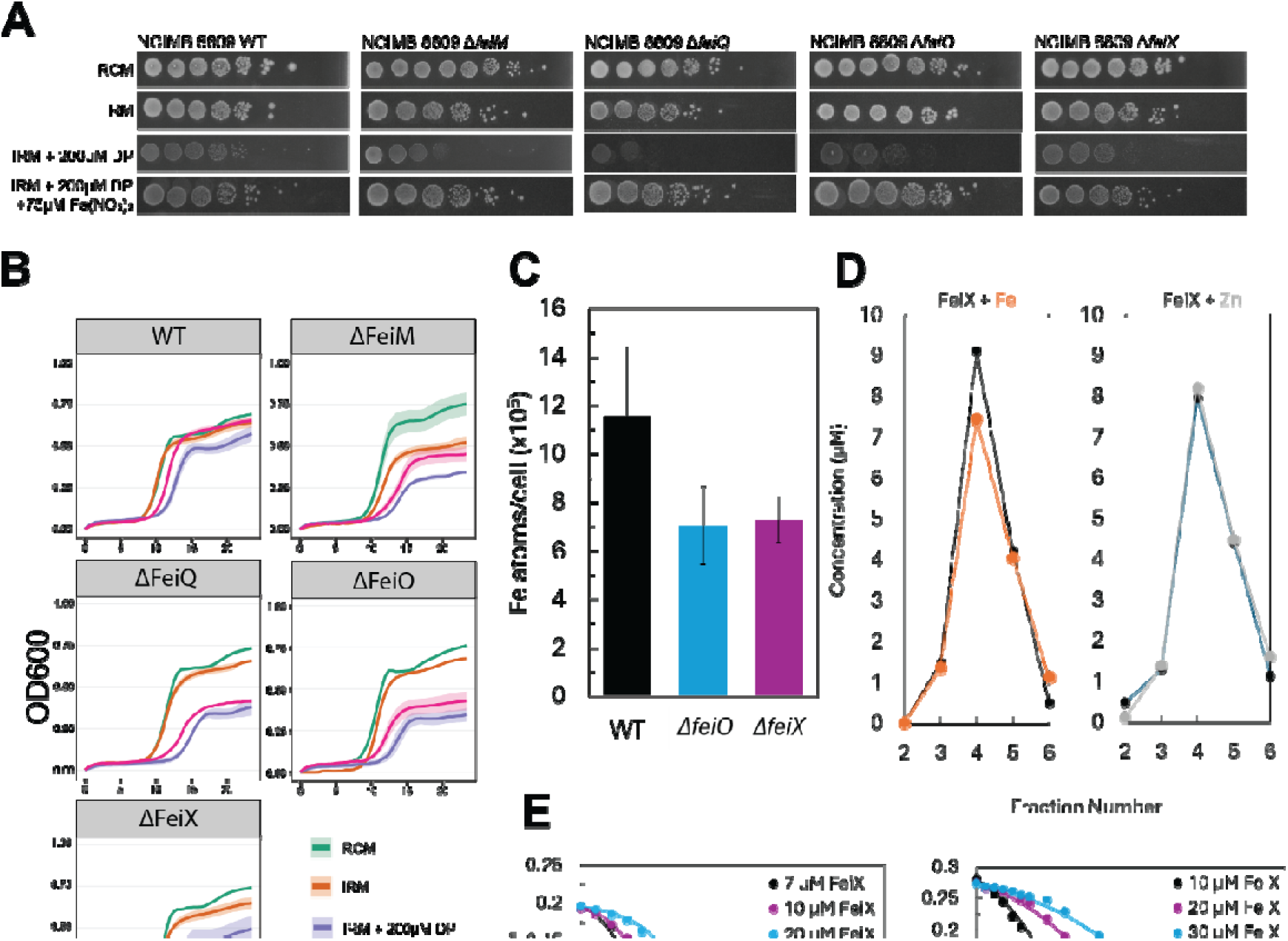
a-e. **The FeiMQOX locus contributes to growth and iron acquisition under iron-limited conditions.** A) Representative 10-fold spot dilution assays of WT, ΔfeiM, ΔfeiQ, ΔfeiO and ΔfeiX strains grown on RCM, IRM, IRM + 200 µM DP, and IRM+ 200 µM DP and 75 µM ferric nitrate grown under anaerobic conditions for 24 hours. B) Growth of WT and FeiMQOX mutant strains in rich control medium (RCM), iron-restricted medium (IRM), IRM supplemented with the iron chelator 2,2′-dipyridyl (DP), or DP supplemented with FeNO3 (DP + Fe). Growth was monitored by measuring OD_600_ over time. Lines represent mean values and shaded regions indicate ± SE from three independent experiments. C) Cellular Iron content on *B. longum* NCIMB 8809 *fei* mutants. Cells were grown for 24 h in IRM + 100 µM bipyridyl. Total iron atoms per cell determined by cell digests as described in the Methods. Error bars indicate standard error of two biological replicates. D) FeiX binds one equivalent of Fe(II) or Zn(II). Protein (•) and metal concentrations (Fe, •, or Zn, •) are indicated. Solid lines are shown for visualisation purposes. E) Fe(II) binding affinity determination in the presence of different FeiX concentrations. Solid lines represent the best global fit to a 1:1 Fe:FeiX binding model. Zn(II) binding affinity determination in the presence of different FeiX concentrations. Solid lines represent fits to a model describing protein competition with MF-2 for 2 equivalents of Zn(II) per monomer.

Supplementation of iron-restricted cultures with ferric iron, partially restored growth of the Fei mutants (Figure 4B and 4C). The physiological rescue demonstrates that the growth defect were associated with iron availability rather than a condition-independent loss of viability. However, growth did not return fully to parental-strain levels in any mutants, suggesting that alternative iron acquisition pathways do not completely compensate for disruption of Fei under the conditions tested.

To determine whether reduced growth was accompanied by altered iron accumulation, metal content was measured by ICP-MS. Under iron-restricted conditions, iron content was reduced by 39.22% and 36.88% in the *feiO* and *feiX* mutants, respectively, relative to the parental strain (Figure 4D). The reduction in cell-associated iron under iron restriction, together with the iron-dependent growth phenotype, supports a direct role for Fei in bacterial iron acquisition. Integrating the transcriptional, genetic, elemental and biochemical findings, we propose that FeiM, FeiQ and FeiO form an ECF-associated membrane complex that supports iron import, while FeiX contributes to iron recognition, binding or delivery to the transport machinery (Figure 4E). These complementary results establish Fei as an iron-responsive acquisition system that supports cellular iron accumulation and growth when iron availability is restricted

## Fei promotes competitive fitness and is associated with microbiota responses to infant iron supplementation

To determine whether Fei contributes to *B. longum* fitness during interspecies competition, we co-cultured parental *B. longum* subsp. *longum* NCIMB 8809 or the corresponding *feiX* mutant with an infant-derived *Enterococcus faecalis* isolate under standard, iron-depleted and iron-supplemented conditions (Figure 5A).

**Figure 5.**
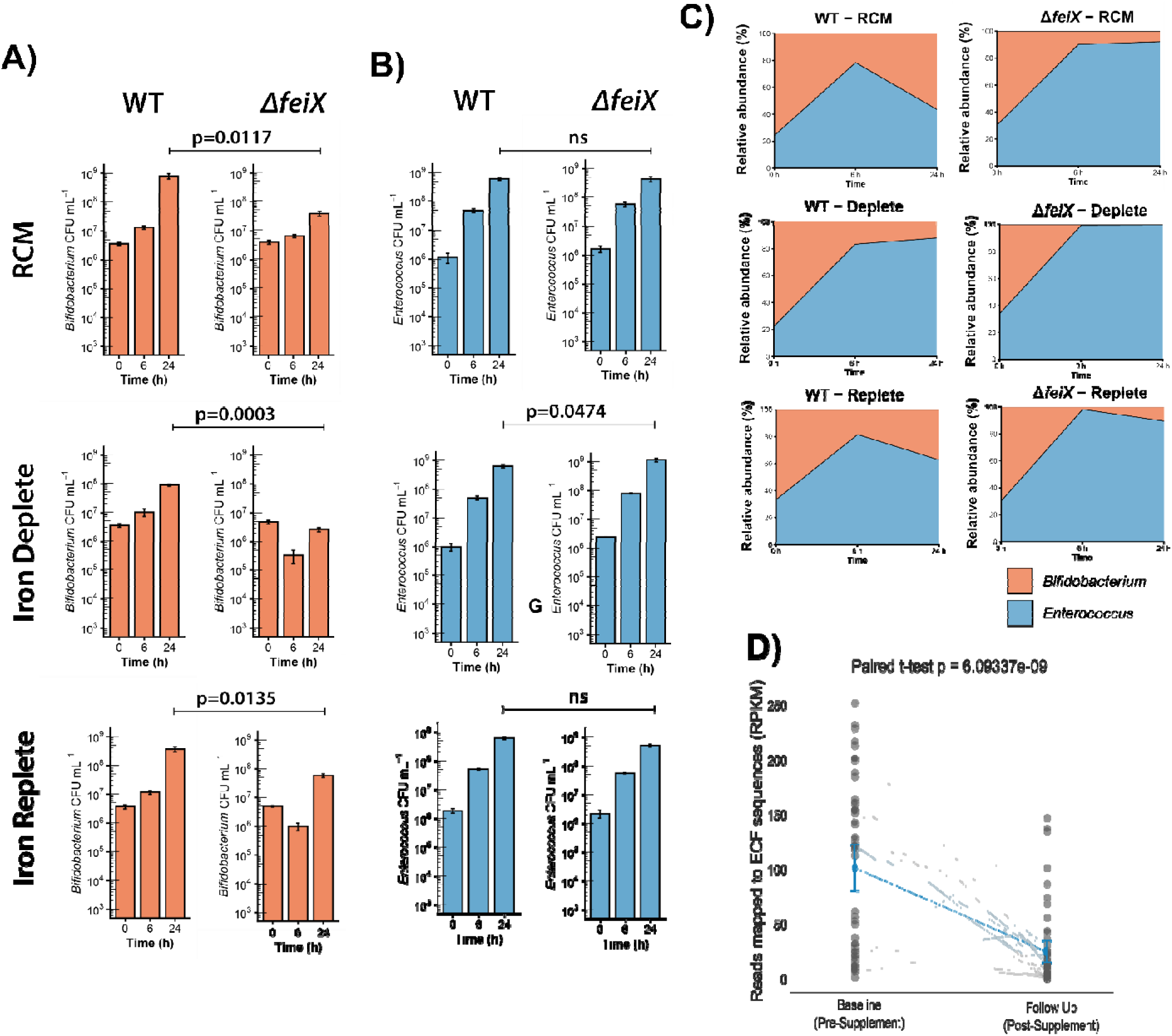
FeiX promotes competitive fitness under iron-limited conditions and is enriched following iron supplementation in vivo. A) Viable counts of *Bifidobacterium longum* subsp. *longum* NCIMB 8809 WT (left) and the Δ*feiX* mutant (right) following co-culture with *Enterococcus faecalis* for 0, 6 and 24 h. Cultures were grown in reinforced clostridial medium (RCM), iron-deplete medium (IRM supplemented with 200 μM 2’-2 Dipyridyl), or iron-replete medium (IRM supplemented with 200 μM 2’-2 Dipyridyl and 75 μM ferric nitrate). Statistical significance was determined between WT and Δ*feiX* strains at the 24 h timepoint using paired t-test. Data represent mean ± SE from 3 independent biological replicates. B) Viable counts of *Enterococcus faecalis* recovered from the same co-culture experiments and conditions described in panel A. Data represent mean ± SE from 3 independent biological replicates. Statistical significance was determined between WT and Δ*feiX* strains at the 24 h timepoint using paired t-test. C) Relative abundance of *B. longum* NCIMB 8809 WT (left) and the Δ*feiX* mutant (right) during co-culture with *E. faecalis*, calculated from viable count data under the iron conditions described in panel A. D) Metagenomic sequence reads mapping to the *feiMQOX* locus in faecal samples from infants enrolled in the BRISC cohort. Read abundance was determined at baseline (pre-ferrous sulphate supplementation) and follow-up (post-supplementation). Statistical significance was assessed using a paired t-test.

After 24 h, recovery of the *feiX* mutant was reduced relative to the parental strain under all three conditions tested. The difference was greatest under iron depletion, where recovery decreased from 8.7x10^7^ CFU/mL for the parental strain to 2.67x10^6^ CFU/mL for the mutant (*p* = 0.0003). Iron supplementation increased recovery of both strains but did not fully restore the competitive fitness of the *feiX* mutant, which remained less abundant than the parental strain at 24 h (*P* = 0.0135; Figure 5B). Notably, disruption of *feiX* did not produce a measurable growth defect in RCM monoculture, whereas recovery of the mutant was reduced during co-culture in the same medium (*P* = 0.0117). Thus, the contribution of FeiX became particularly apparent during interspecies competition.

Loss of FeiX also altered the composition of the two-member community. Under iron depletion conditions, *B. longum* represented 12.04% of the recoverable population in parental-strain co-cultures but only 0.23% in mutant co-cultures. Iron supplementation partially restored this defect, although *B. longum* still represented only 10.45% of the population in mutant co-cultures compared with 37.3% in the parental-strain co-cultures (Figure 5C). Recovery of *E. faecalis* was correspondingly greater in iron-depleted co-cultures containing the *feiX* mutant than in those containing parental *B. longum* (1.17x10^9^ versus 6.33x10^8^ CFU/mL; *p* = 0.0473). These findings demonstrate that FeiX supports *B. longum* fitness during competition, particularly when iron availability is restricted.

We next examined the relevance of Fei within an iron-variable infant gut ecosystem by reanalysing shotgun metagenomic data from an iron-supplementation study in rural Bangladesh (9). The analysis included paired baseline and post-intervention samples from 36 infants receiving ferrous sulphate (Figure 5D). Reads mapping to the *feiMQOX* locus were detected in 51 infants at both baseline and following supplementation. Paired analysis identified a reduction in normalised Fei abundance following iron supplementation, from an average of 101.55 RPKM at baseline to 25.02 RPKM after treatment (estimated average change, -76.53 RPKM; 95% confidence interval, -98.51 RPKM to -54.65; *p* = 6.09337e-09; Figure 5E). To establish the taxonomic and genomic context of these sequences, we reconstructed metagenome-assembled genomes from the wider cohort. We recovered 25 high-quality MAGs classified as *B. longum*, and the complete *feiMQOX* locus was detected in 15 of these genomes. By contrast, we recovered 103 *B. infantis* MAGs, none of which contained the full *feiMQOX* locus. (Supplementary Figure 2). Together, these data show that Fei supports *B. longum* during microbial competition and that the abundance of Fei-encoding bacteria changes following iron supplementation in infants. The direction of this change is consistent with Fei providing its greatest relative advantage under iron-restricted conditions, although the cohort analysis remains associative and does not establish Fei-dependent persistence *in vivo*.

## Discussion

The ability to acquire limiting nutrients is fundamental to microbial persistence, yet this aspect of gut microbiota ecology has been studied predominantly in pathogens and Gram-negative bacteria. Our findings identify Fei as an iron-responsive acquisition system that links micronutrient availability to the competitive fitness of *B. longum* subsp. *longum*.

Integration of transcriptional, genetic, biochemical and elemental analyses supports a role for Fei in iron acquisition, while the competition experiments show that its ecological importance extends beyond growth in monoculture. The uneven distribution of *feiMQOX* across bifidobacterial lineages, together with its presence in infant gut metagenomes and changing abundance following iron supplementation, further suggests that iron acquisition represents a species-level trait through which bifidobacteria respond to nutritional and microbial pressures in the gut.

Iron acquisition is often considered in the context of pathogens, where siderophores, haem uptake and ferrous or ferric iron transport systems contribute to colonisation and virulence (54–56). However, the same resource constraints operate across the wider microbiota, particularly in the colon, where iron is poorly soluble, frequently bound within host, dietary or microbial complexes, and subject to intense competition (57). Therefore, beneficial anaerobic bacteria must balance their iron requirements against investment in acquisition systems whose value will depend on the local chemical and microbial environment. Indeed, the marked variation in iron-limited growth across the bifidobacterial strains examined supports this concept, as iron tolerance was not uniform, rather multiple *B. longum* subsp. *longum* strains retained growth under conditions that strongly restricted other bifidobacteria. The conservation of Fei within this lineage highlights a potential mechanistic basis for part of this phenotype, while the residual growth of Fei mutants indicates that Fei likely acts within a broader network of iron acquisition and metabolic adaptation.

The association of Fei with *B. longum* subsp. *longum* may be particularly relevant to the ecological range of this subspecies. Unlike bifidobacterial lineages most strongly associated with exclusive milk feeding, *B. longum* subsp. *longum* is detected from early infancy through weaning and into adulthood, and individual strains can persist across substantial changes in diet and microbiota composition (58). This broader and more sustained niche exposes *B. longum* subsp. *longum* to changing iron availability, carbon sources, pH and microbial competitors. Therefore, a high-affinity iron acquisition system could offer additional resilience across successive gut environments rather than adaptation to only one developmental stage. This interpretation is consistent with the strong conservation of the complete *feiMQOX* locus within *B. longum* subsp. *longum*, but its absence or incomplete representation in several closely related lineages such as *B. longum* subsp. *infantis*. Importantly, Fei is unlikely to explain the ecological distribution of *B. longum* subsp. *longum* alone. Carbohydrate utilisation, stress tolerance, cell-surface structures and interactions with the host will act alongside micronutrient acquisition to determine strain persistence (59–61). Fei should therefore be viewed as one component of a wider suite of traits that enables this subspecies to remain competitive under changing gut environments and ecosystems.

Fei also represents a previously unknown solution to bacterial iron acquisition. ECF transporters are generally associated with high-affinity uptake of vitamins and selected (cobalt and nickel) transition-metal substrates, whereas iron acquisition more commonly involves siderophores, haem receptors, ferric-binding proteins or dedicated ferrous iron transporters (54–56). Induction of the complete *feiMQOX* locus during iron depletion, reversal towards controlled expression following iron addition, reduced cell-associated iron in Fei mutants and partial physiological rescue by iron collectively connect this ECF-associated system to iron homeostasis. FeiX provides an additional distinguishing feature as its predicted structural relationship to a known metal-binding protein and its affinity for Fe(II) suggest that it could contribute to intracellular iron handling. The current data do not establish whether FeiX receives iron directly from the membrane complex or functions after import into the cytosol. Nevertheless, the presence of FeiX alongside the core ECF-associated components may help explain how a transporter family principally associated with other micronutrients has been adapted for iron acquisition. The recurrent absence of FeiX from some otherwise related loci further raises the possibility that complete and partial Fei systems differ functionally, rather than representing equivalent variants of the same pathway.

The ecological value of Fei became most apparent in the presence of a competitor. Loss of FeiX produced a substantially greater fitness defect during co-culture with *E. faecalis* than during monoculture, including in conditions in which iron availability was sufficient to support apparently normal monoculture growth. Thus, the amount of iron required to sustain growth in isolation may differ from that required to remain competitive within a community. This distinction is likely important because nutrient requirements defined in monoculture can underestimate the selective value of high-affinity acquisition pathways in the dynamic gut environment. Increased recovery of *E. faecalis* when competing with the *feiX* mutant under iron limitation is consistent with competition for a restricted resource, although indirect effects of Fei disruption on *B. longum* metabolism cannot be excluded. More broadly, these findings suggest that Fei may influence community assembly without directly antagonising other organisms; allowing *B. longum* to retain a larger share of a limiting micronutrient, with Fei supporting bifidobacterial abundance and indirectly constraining organisms occupying overlapping nutritional niches. Whether this extends to efficient iron scavengers such as Enterobacteriaceae, or operates within more complex infant communities, is an important question for future work.

The infant cohort analysis helps place this mechanism within a clinically relevant context in which iron availability is deliberately altered due to underlying IDA. A reduction in normalised Fei abundance following supplementation is consistent with Fei providing its greatest relative advantage when iron is scarce. Iron supplementation may reduce the benefit of maintaining high-affinity iron acquisition, while simultaneously increasing iron availability to organisms that compete effectively under iron-replete conditions. However, due to the lower metagenomic coverage data, we cannot distinguish altered representation of Fei-positive *B. longum* strains from an overall change in *B. longum* abundance or wider community composition. The cohort findings therefore support the ecological relevance of Fei but do not establish that Fei directly drives persistence within the infant gut. Integrating Fei abundance with total *B. longum* coverage, baseline iron status and longitudinal strain-resolved genomes will be important for determining whether the locus is specifically associated with persistence during iron deficiency or with responses to supplementation. This may also help explain why microbiota responses to iron interventions vary substantially among infant populations, since the outcome may depend partly on the strain-level iron acquisition traits already present within the community.

These findings could inform microbiota-directed approaches designed to accompany, rather than replace, clinically necessary iron interventions. Fei-positive *B. longum* subsp. *longum* strains may be promising next-generation probiotic candidates because they combine relevance to early life with the capacity for longer-term persistence across weaning and into adulthood. Their ability to remain competitive under changing iron conditions may be particularly valuable where supplementation modifies the balance between beneficial commensals and efficient iron-scavenging pathobionts. However, Fei carriage should be treated as one functional selection criterion rather than a standalone probiotic marker. Candidate strains would also require appropriate carbohydrate-utilisation capacity, safety, genomic stability and evidence of fitness within representative microbial communities. Strains isolated from the intended geographical populations, or rigorously validated within those populations, should be prioritised because bifidobacterial lineages, diet, iron status and background microbiota differ between settings (62,63).

There are several limitations which could be explored in future work. The iron-restricted media and chelation conditions used here cannot reproduce the chemical and spatial complexity of the intestinal environment, and insertional mutagenesis may have affected expression of downstream genes. Genetic complementation and direct measurements of iron-transport kinetics will therefore be important for resolving the contributions of individual Fei components. Similarly, ICP-MS measures total cell-associated iron and not its oxidation state or localisation. Defining the physiological Fei substrate and the precise function of FeiX will be central to establishing the molecular mechanism. Finally, the two-member competition model and associative cohort analysis could be extended using strain-resolved longitudinal data, representative infant microbial communities and *in vivo* models of iron deficiency and supplementation. These approaches may help explore how Fei supports long-term colonisation and whether its effects extend beyond individual competitors to wider ecosystem structure.

In conclusion, our findings identify Fei as a lineage-associated iron acquisition system that allows *B. longum* subsp. *longum* to translate access to a limiting micronutrient into competitive fitness. This capacity may be particularly important for a bifidobacterial lineage that occupies the gut across changing nutritional and developmental states. By connecting molecular iron acquisition with microbial competition and responses to infant iron supplementation, this study highlights micronutrient acquisition as an underappreciated determinant of beneficial bacterial persistence and provides a foundation for function-led selection of microbiota-directed interventions.

## Acknowledgements

L.J.H. is funded by a Wellcome Trust Investigator Award no. 220876/Z/20/Z, and a Biotechnology and Biological Sciences Research Council (BBSRC) Institute Strategic Programme Food Microbiome and Health BB/X011054/1 and its constituent project BBS/E/F/000PR13631. L.J.H. L.A. are supported by a BBSRC Responsive Mode Grant BB/Y011619/1. P.T.C. and J.A. are supported by a BBSRC Responsive Mode Grant BB/Y009282/1. D.K.J.S. is supported by a Colin Charles Frederick Blake University of Birmingham PhD Studentship with L.J.H. as supervisor.

## Author contributions

Conceptualisation: L.A., P.T.C. and L.J.H.; Methodology: L.A., C.Z.S., A.S. and J.A.; Software: H.T. and M.K.; Validation: L.A. and J.A.; Formal analysis: L.A., S.C., J.A.,-C.Z.S. and M.K.; Investigation: L.A., C.Z.S., J.A., and A.S.; Resources: C.C., L.F. and D.v.S.; Writing – original draft: L.A., P.T.C. and L.J.H.; Writing – review and editing: L.A, P.TC, L.J.H.; Visualisation: L.A., P.T.C. and M.K.; Supervision: L.J.H.; Funding acquisition: P.T.C. and L.J.H.

## Data availability

RNA-sequencing data generated in this study have been deposited in National Centre for Biotechnology Information Sequence Read Archive (SRA) under accession number PRJNA1559618. Sequencing readds generated for mutant validation have been deposited in SRA under accession numbers PRJNA1559618. The publicly available BRISC metagenomic data analysed in this study are available from NCBI under accession number PRJNA1081952.

