## Supplementary Information for "An iron acquisition system promotes *Bifidobacterium longum* fitness in competitive early life gut communities"

Supplementary Data

**Table 1 – Strains used in this study**

Supplementary Table 1: List of Bacterial Strains used within this study

| Strain | Reference/Source |
| --- | --- |
| Bifidobacterium longum subsp Longum LH277 | SAMEA4838171 |
| Bifidobacterium longum subsp. longum NCIMB 8809 | CP011964 |
| *Escherichia coli* strain EC101 | Gift from Douwe van Sinderen |
| Bifidobacterium longum subsp. longum NCIMB 8809 Δ*feiM* | This Study |
| Bifidobacterium longum subsp. longum NCIMB 8809 Δ*feiQ* | This Study |
| Bifidobacterium longum subsp. longum NCIMB 8809 Δ*feiO* | This Study |
| Bifidobacterium longum subsp. longum NCIMB 8809 Δ*feiX* | This Study |
| Bifidobacterium longum subsp. infantis YCFA59 | This Study |
| Bifidobacterium longum subsp. infantis B53 | This Study |
| Bifidobacterium longum subsp. infantis B55 | This Study |
| Bifidobacterium longum subsp. infantis DSM20088 | CP001095.1 |
| Bifidobacterium longum subsp. infantis MUP21 | This Study |
| Bifidobacterium mellis LH898 | SAMN45905979 |
| Bifidobacterium psychraerophilum LHZ012 | SAMN45905982 |
| Bifidobacterium aesculapii LH461 | SAMN45905937 |
| Bifidobacterium callitrichidarum LH423 | SAMN45905921 |
| Bifidobacterium reuteri LH307 | SAMN45905894 |
| Bifidobacterium breve TOS12 | This study |
| Bifidobacterium breve UCC2003 | CP000303 |
| Bifidobacterium breve LH21 | This study |
| Bifidobacterium breve MUP22 | This study |
| Bifidobacterium breve DSM20013 | AP012324 |
| Bifidobacterium longum subsp. longum E2T2-1 | This Study |
| Bifidobacterium longum subsp. longum DSM20219 | ASM19655v1 |
| Bifidobacterium longum subsp. longum TOSE4 | This Study |

Table 2 – Primers used in this study

Supplementary table 2 – Primers and Plasmids used in this study

| Primer/Plasmid | Reference/Source | Purpose |
| --- | --- | --- |
| pFREM2 | Gift from Prof. Douwe van Sinderen | Cloning Plasmid with B.*longum* restriction motif’s removed |
| EcfORF_FW_XbaI | gagatcTCTAGAAACTGGAGGTCTCCGACCTC | Amplify 477bp fragment, add XbaI sites |
| EcfORF_RV_NcoI | gagatcCCATGGACGAATACGCCGTCGATGTA | Amplify 477bp fragment, add NcoI sites |
| EcfS_FW_XbaI | gagatcTCTAGAAGCAGGCGGTGCTCGCCAA | Amplify 411bp fragment, add XbaI sites |
| EcfS_RV_NcoI | gagatcCCATGGTGGCGTACTGGAAATGCAGG | Amplify 411bp fragment, add NcoI sites |
| EcfA_FW_XbaI | gagatcTCTAGAGGTATTGGATGAGCCGACC | Amplify 516bp fragment, add XbaI sites |
| EcfA_RV_HindIII | gagatcAAGCTTGCTCGATGCGGCCCGAAACA | Amplify 516bp fragment, add HindIII sites |
| EcfT_FW_XbaI | gagatcTCTAGAGACTGCCGTGGCTGCATGTG | Amplify 485bp fragment, add XbaI sites |
| EcfT_RV_NcoI | gagatcCCATGGTCAGATCACCCCCGCGATAT | Amplify 485bp fragment, add NcoI sites |
| CK_FeiM_FW | ATCACCGTCATGGGCGTGATCAT | Check FeiM insertion |
| CK_FeiQ_FW | TTCGCAATGCGATGGCGTTGCG | Check FeiQ insertion |
| CK_FeiO_FW | TTACGAGGCCGATGAATTCCTGT | Check FeiO insertion |
| CK_FeiX_FW | ACGACACCAGCCTCGACACCTA | Check FeiX insertion |
| Ck_pFrem_RV | TGCTTCCGTACTACGACCCC | Check Insertional mutants |

**Table 3 – Sequence data used for Bifidobacterium phylogenetic tree in Figure 3**

***See Attached File***

**Figure 1 – BLAST of FeiMQOX region**

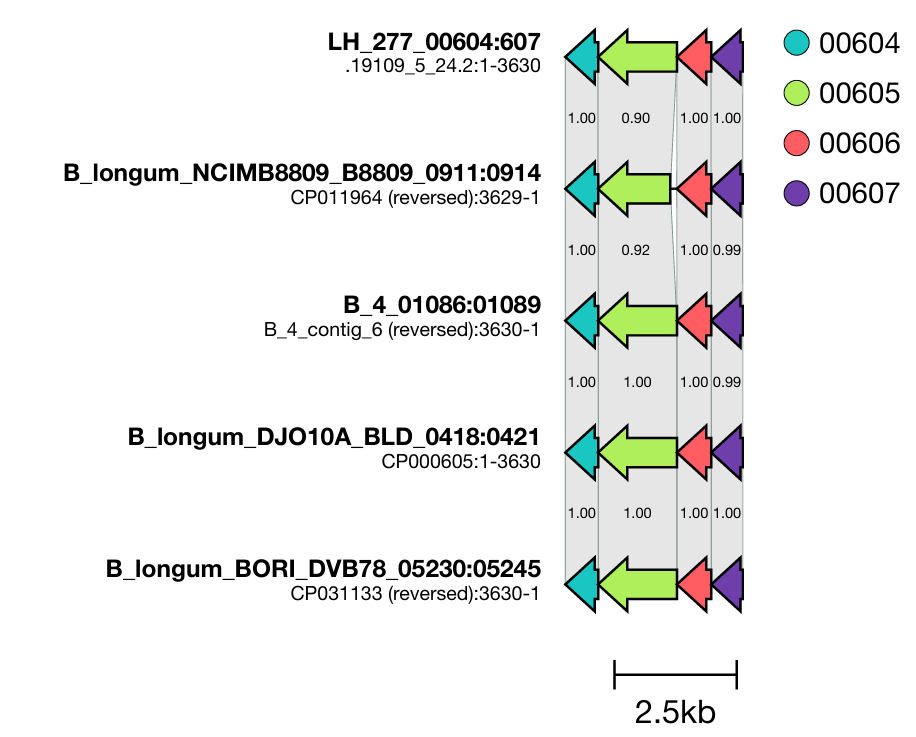

**Supplementary Figure 1** – BLAST alignment comparisons of ECF regions from selected genomes. FeiM = 607, FeiQ=606, FeiO=605 and FeiX=604

Figure 2 – MAGs from BRISC study

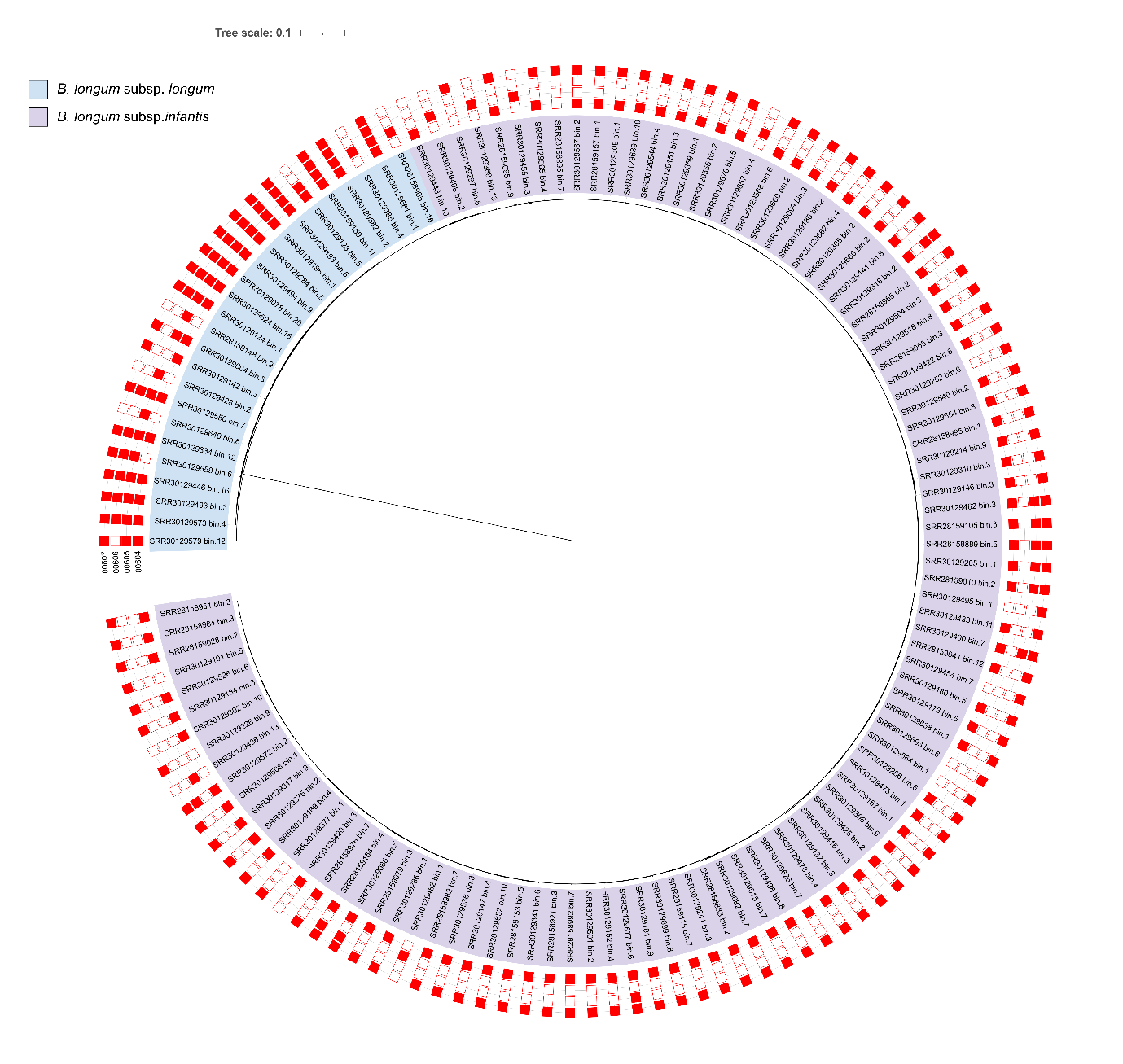

Supplementary Figure 2 – *Bifidobacterium longum metagenome assembled genomes from the BRISC cohort study. Bifidobacterium longum subsp. longum = blue, Bifidobacterium longum subsp. infantis = purple. Presence or absence of FeiMQOX, presence = Red, absence = white.* FeiM = 607, FeiQ=606, FeiO=605 and FeiX=604
